# Retinal direction selectivity in the absence of asymmetric starburst amacrine cell responses

**DOI:** 10.1101/428532

**Authors:** Laura Hanson, Santhosh Sethuramanujam, Geoff deRosenroll, Gautam B. Awatramani

## Abstract

In the mammalian retina, asymmetric inhibitory signals arising from the direction-selective dendrites of GABAergic/cholinergic starburst amacrine cells are thought to be crucial for originating direction selectivity. Contrary to this notion, however, we found that direction selectivity in downstream ganglion cells remains remarkably unaffected when starburst output is rendered non-directional (using a novel strategy combining a conditional GABA_A_ α2 receptor knockout mouse with optogenetics). We show that temporal asymmetries between excitation/inhibition, arising from the differential connectivity patterns of starburst cholinergic and GABAergic synapses to ganglion cells, form the basis for a parallel mechanism generating direction selectivity. We further demonstrate that these distinct mechanisms work in a coordinated way to refine direction selectivity as the stimulus crosses the ganglion cell’s receptive field. Thus, precise spatiotemporal patterns of inhibition and excitation that shape directional responses in ganglion cells are shaped by two ‘core’ mechanisms, both arising from distinct specializations of the starburst network.

## Introduction

Understanding how neural circuits in the brain compute information not only requires determining how individual inhibitory and excitatory elements of circuits are wired together, but also a detailed knowledge of their functional interactions. Recent advances in optogenetic techniques and mouse genetics now offer ways to specifically probe the functional properties of neural circuits with unprecedented specificity. Perhaps one of the most heavily interrogated circuits in the mouse brain is one in the retina that is involved in coding direction (reviewed by Mauss et al., 2017; Vaney et al., 2012). Here, direction is encoded by specialized direction-selective (DS) ganglion cells (DSGCs), which respond robustly to objects moving in a ‘preferred’ direction but not in the opposite or ‘null’ direction (Barlow and Levick, 1965), which we now know relies on the coordination of three transmitter systems: glutamate, GABA and acetylcholine. However, despite the in-depth functional and anatomical characterization of this circuit, the precise spatiotemporal dynamics of each system still remains unclear and thus how directional selectivity emerges in ganglion cells remains to be fully elucidated.

In prevailing models, direction selectivity is shaped by the average excitation/inhibition (E/I) ratio (reviewed by Mauss et al., 2017; Vaney et al., 2012). Null-direction motion evokes large amplitude inhibitory postsynaptic currents (IPSCs) in DSGCs (Fried et al., 2002; Taylor and Vaney, 2002), which effectively counteract coincident excitation, mediated by glutamatergic and cholinergic input. During preferred-direction motion, the inhibitory charge is significantly weaker, enabling excitatory inputs to drive robust spiking responses in DSGCs. Corroborative evidence from a number of seminal studies indicates that DS inhibition is largely mediated by direction-selective dendrites of starburst amacrine cells (Briggman et al., 2011; Ding et al., 2016; Euler et al., 2002; Fried et al., 2002; Lee et al., 2010; Taylor and Vaney, 2002; Yonehara et al., 2011; Yoshida et al., 2001) (but see Pei et al., 2015). However, the role of direction selectivity in starburst cells has been difficult to confirm as the underlying mechanisms are still being evaluated and to date there has been no easy way to selectively abolish it. Nevertheless, most studies consider changes in E/I amplitude as the major factor that underlies direction selectivity in ganglion cells (Yonehara et al., 2016; Morrie and Feller, 2018; Pei et al., 2015; Koren et al., 2017, Poleg-Polsky and Diamond 2016; Chen et al., 2016)

Before the discovery of direction selectivity in starburst dendrites, early theoretical studies postulated that direction selectivity emerged from a temporal asymmetry between inhibition and excitation, in which excitation activates first during preferred-direction motion, and inhibition activates first during null-direction motion (Koch et al., 1982; Taylor et al., 2000; Torre and Poggio, 1978). Several lines of recent evidence suggests that such E/I offsets could be the natural outcome of the asymmetrical wiring of the starbursts to DSGCs. Specifically, elegant starburst/DSGC paired recordings directly demonstrate that the cholinergic inputs arise symmetrically from starbursts surrounding the DSGCs, while GABAergic inputs arise only from starbursts with somas displaced to the null-side of the DSGC’s receptive field (i.e. the side from which null-stimuli enter the receptive field; Lee et al., 2010; Fried et al; Yonehara et al., 2015; Chen et al.,2017; Brombas et al., 2017). Indeed, this arrangement produces the required E/I timing differences in DSGCs, when stimulated naturally with moving stimuli (Sethuramanujam et al., 2016).

While temporal E/I asymmetries have often been noted in the literature, their impact has been difficult to assess, as they are always associated with changes in E/I amplitude ratio (Fried et al., 2005; Kostadinov and Sanes, 2015; Pei et al., 2015; Sethuramanujam et al., 2016; Taylor and Vaney, 2002). Further, blocking cholinergic inputs, which is expected to decrease E/I timing differences arising from the abovementioned specific wiring of starburst ACh/GABA synapses, does not appear to affect the ability of DSGCs cells to encode direction (Ariel and Daw, 1982; Kittila and Massey et al., 1997; Taylor and Smith, 2012). Additionally, modeling studies suggest that E/I offsets play a negligible role in this computation in the context of the E/I amplitude modulation (Schachter et al., 2010).

A major caveat in the interpretation of results from previous studies evaluating the role of ACh and E/I temporal asymmetries in direction selectivity, however, is that they assume a single “core” mechanism underlies direction selectivity. Here, we found that blocking starburst DS, using a novel combination of optogenetics/mouse KO technology/pharmacology, had a surprisingly weak effect on direction selectivity in ganglion cells, demonstrating the existence of a second DS mechanism. Analysis of the synaptic inputs shows that E/I timing differences arising from the specific wiring of starburst to DSGCs serves as the substrate for this parallel mechanism. Interestingly, while both changes in E/I ratios or timing differences are sufficient to drive robust DS responses in ganglion cells, they each contribute to distinct phases of the DSGC response, ensuring that direction is computed rapidly, with high fidelity.

## Results and Discussion

### Rendering starburst dendrites non-directional

Here, we first sought to develop ways to block direction selectivity in starburst dendrites in order to test its role in originating direction selectivity in the retina. Recent studies have put forth two distinct models relying on the properties of inhibitory (Ding et al., 2016; Lee and Zhou, 2006; Munch and Werblin, 2006) or excitatory networks (Fransen and Borghuis, 2017; Kim et al., 2014) that could potentially underlie direction selectivity in starburst dendrites. But when experimentally tested, neither of these mechanisms appears to be required. For example, the inhibitory network mechanism is based on mutual inhibition between anti-parallel starburst dendrites (Ding et al., 2016; Lee and Zhou, 2006; Munch and Werblin, 2006), but selectively eliminating it using the GABA_A_ α2 receptor KO mouse line (*Gabra2* KO) leaves starburst DS largely intact (Chen et al., 2016). Consistent with this notion, we found the average direction selectivity index (DSI; see methods) of IPSCs measured in DSGCs in *Gabra2* KO mice, was comparable to that measured in wild type DSGCs (Wt = 0.33 ± 0.019, n = 6; KO = 0.28 ± 0.022; n = 6; Fig. 1a, b,e,f). Similarly, the competing excitatory network model, in which the selective wiring of temporally distinct bipolar cells along the proximal-distal axis of the starburst dendrite is proposed to originate direction selectivity (Fransen and Borghuis, 2017; Kim et al., 2014), has also been called into question as the direct stimulation of the channelrhosdopsin2 (ChR2)-expressing starburst network in relative isolation (i.e. with glutamate inputs blocked) generates robust DS starburst output (Sethuramanujam et al., 2016) (Fig. 1c,e,f).

**Fig. 1:**
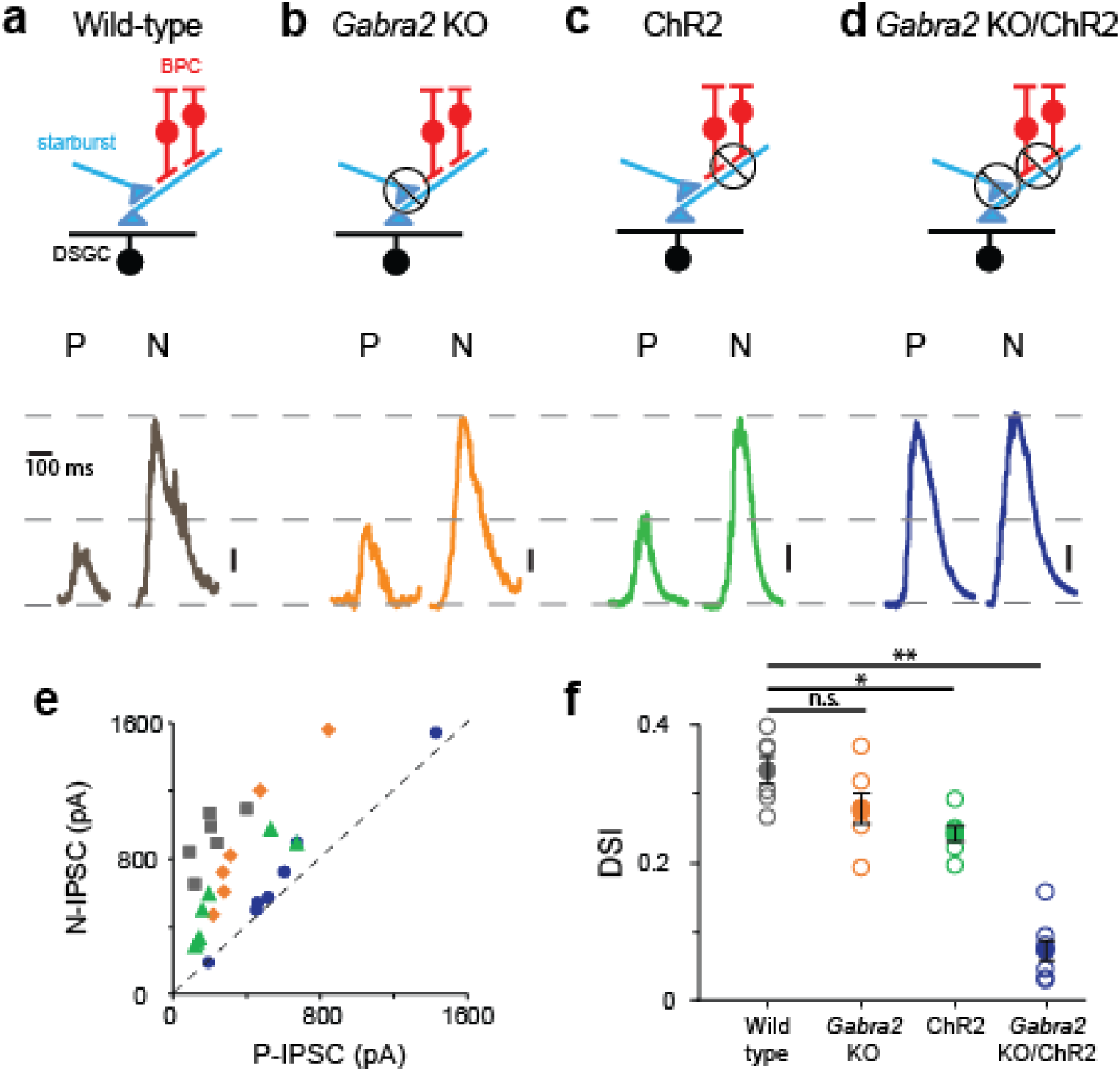
Direction selectivity in starburst dendrites relies on excitatory and inhibitory network mechanisms. **a-d**, IPSCs recorded in voltage-clamped DSGCs (V_HOLD_ ~0 mV) in a variety of mouse lines, as indicated in the top panels. Responses are shown for stimulus motion in the DSGC’s preferred (P) or null-direction (N) under conditions in which **a**, all synaptic inputs are intact; **b**, mutual inhibition between starbursts is selectively disrupted using the conditional *Gabra2* KO; **c**, bipolar cell inputs are blocked and ChR2-expressing starbursts are directly stimulated; **d**, both mutual inhibition and bipolar cell inputs are blocked. Vertical scale bar = 100 pA (**a-c**) or 200pA (**d**). **e**, The peak amplitudes of the IPSCs evoked during preferred and null motion are plotted against each other for the conditions noted in **a-d**. Reference line (dashed line; slope = 1) is indicated. n=6 for wild type, *Gabra2* KO and ChR2; n=7 for *Gabra2* KO/ChR2. **f**, The average direction-selectivity index (DSI) computed from the vector sum of the peak amplitude of IPSCs evoked by stimuli moving in eight directions, for the conditions in **a-d** (See Methods for DSI calculation; DSI = 0 indicates non-directional responses; DSI = 1 indicates that responses were evoked in only one direction). Pooled data are represented as mean ± SEM (Solid circles), while single cell responses are denoted by open circles. Statistical significance was estimated by unpaired t-tests, where * indicates p < 0.05 and ** indicates p < 0.001.

Remarkably, combining these two approaches to block both inhibitory and excitatory network DS mechanisms, led to the near-complete block of the asymmetry in responses mediated by starburst cells, i.e the amplitudes of inhibitory inputs evoked by optogenetic stimulation of starbursts in the *Gabra2* KO background were nearly equal for preferred and null-direction motion (DSI = 0.07 ± 0.02, n=7; Fig. 1d-f). The ability to block direction selectivity in starbursts, while leaving its output relatively intact, provides for the first time, direct evidence for the mechanisms generating it. The requirement to block both the excitatory (Fransen and Borghuis, 2017; Kim et al., 2014) and inhibitory network mechanisms (Ding et al., 2016; Lee and Zhou, 2006; Munch and Werblin, 2006) suggest that they work in parallel to shape DS responses in starburst dendrites.

### Retinal direction selectivity in the absence of asymmetric starburst amacrine cell responses

Abolishing the directional properties of starbursts is expected to supress the output of DSGCs, as in these conditions DSGCs would receive strong inhibition in all directions. However, contrary to this notion, we found that DSGCs in *Gabra2* KO mice continued to exhibit robust spiking responses when starburst output was rendered non-DS (Fig. 2a). Surprisingly, these spiking responses were robustly tuned for direction similar to control conditions (Fig. 2b-d). Both the DSI and the direction encoded were relatively constant across a range of velocities (Fig. 2e). While the direction encoded was the same under conditions in which starburst output was DS or non-DS (Fig. 2a), we did observe a significant reduction in the duration of the DSGC’s spiking response (Fig. 2a). Since starbursts are critically required for DS computation (Vlasits et al., 2014; Yoshida et al., 2001), it follows that they must utilize an alternate mechanism to confer direction selectivity upon DSGCs in the absence of amplitude modulation of inhibitory inputs. In addition, the finding that the direction encoded by the DSGC remains unchanged—when starburst output is rendered non-DS—indicates that the two DS mechanisms must be well-aligned (Fig. 2a-c).

**Fig. 2:**
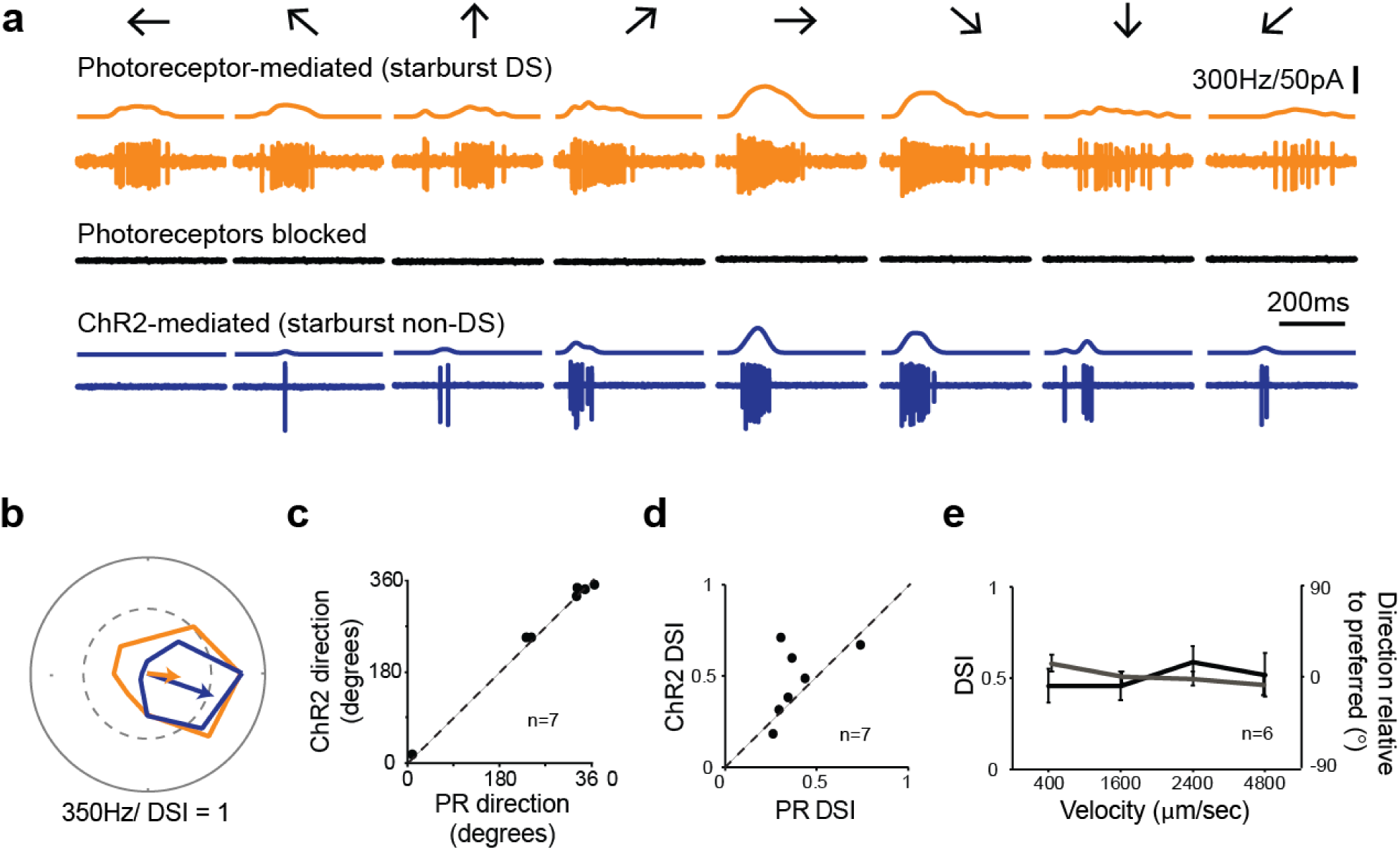
Retinal direction selectivity in the absence of asymmetric starburst amacrine cell responses. **a**, Spiking responses from a DSGC in the *Gabra2* KO/ChR2 mouse line, during photoreceptor mediated stimulation (*top*), when photoreceptor synapses are blocked pharmacologically (50 μM DL-AP4, 10 μM UBP310 and 20 μM CNQX (*middle*)), and when stimulus intensity is increased to directly activate starbursts using ChR2 (*bottom*). This enables a direct comparison of direction selectivity in a DSGC under conditions in which starbursts output is DS (*top*) or non-DS (*bottom*). Stimuli were moved in eight directions indicated by the arrows at a velocity of 1.6 mm/s. The smooth traces indicate the average spike rate estimated by low-pass filtering the spike train via convolution with a Gaussian kernel (σ = 25 ms). **b**, Polar plots of the peak spike rates for the responses shown in **a**. The arrow indicates the DSGC’s preferred direction, scaled to the DSI. **c-d**, A comparison of the preferred direction (**c**) and DSI (**d**) of the responses evoked during ChR2 stimulation and during intact photoreceptor stimulation in the *Gabra2* KO/ChR2 mouse line. **e**, DSI (black) and direction relative to the preferred direction (grey) is plotted as a function of stimulus velocity for ChR2-evoked responses measured from DSGCs in the *Gabra2* KO/ChR2 mouse line. Data are represented as mean ± SEM.

When we examined the onset latencies for GABA and ACh in the *Gabra2* KO, we found E/I offsets were exquisitely tuned for direction (Fig. 3a-c; in the absence of any changes in the E/I amplitudes). The magnitude of the offsets provided a good indication of the DSGC’s preferred-direction when compared to its spiking responses measured prior to the voltage-clamp experiments (Fig. 3b, c). These results provided strong evidence that temporal asymmetries in E/I alone can drive direction selectivity in ganglion cells.

**Fig. 3:**
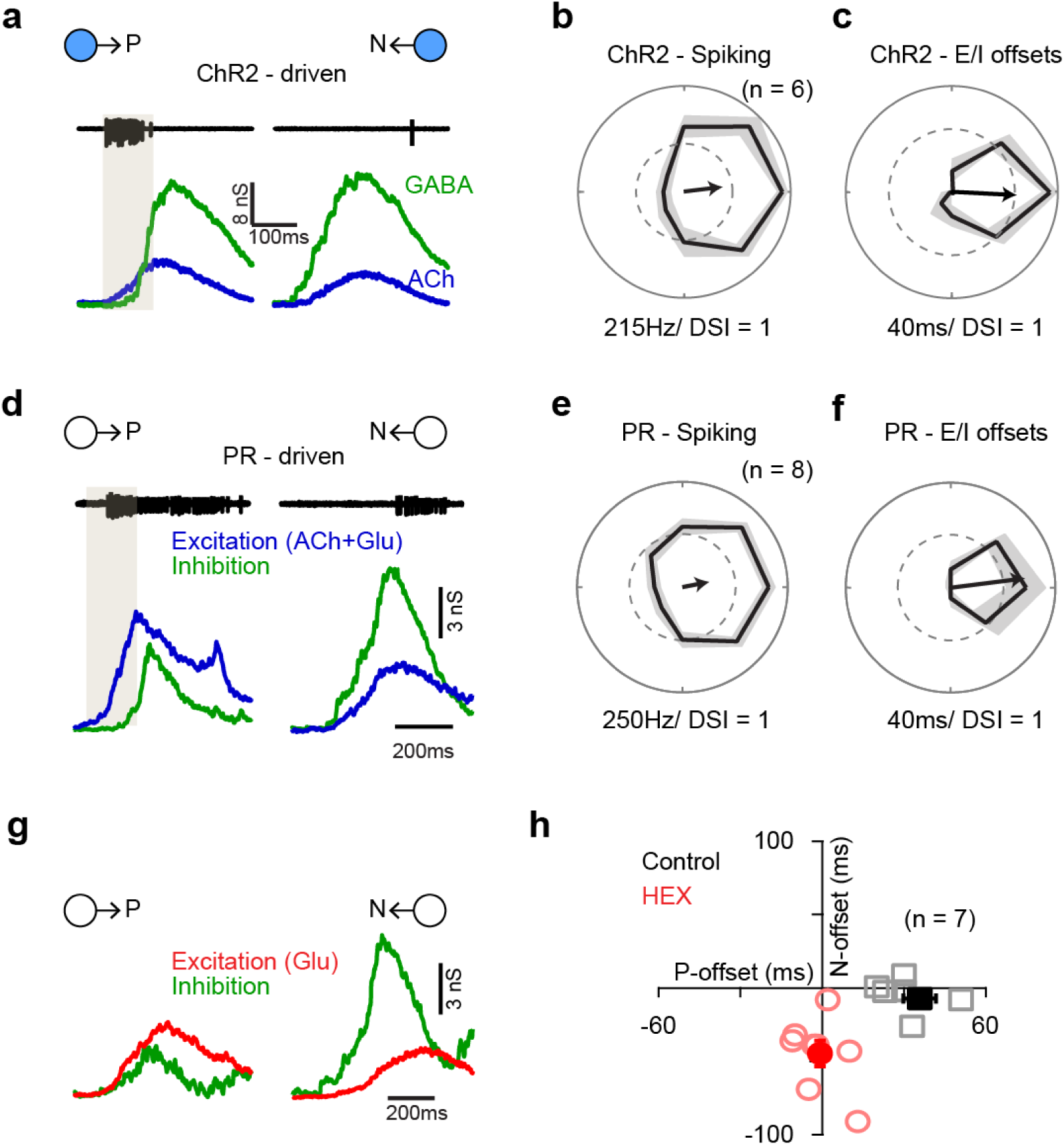
Differential functional wiring of cholinergic and GABAergic starburst synapses shape directionally tuned E/I offsets. **a**, Spiking, EPSCs and IPSCs recorded in the same DSGC evoked by stimulating ChR2 expressing starbursts in the *Gabra2* KO/ChR2 mouse line. Shaded box indicates the E/I offset window in which a large fraction of the spiking occurs. **b-c**, Polar plots of the average peak firing rates (**b**) and E/I offsets (**c**) measured under the conditions described in a, across 6 cells (See Methods for E/I offset estimation). Data are represented as mean ± SEM. **d-f**, Similar to **a-c**, but for responses measured under physiological conditions i.e. responses driven by photoreceptors. Data are represented as mean ± SEM. **g**, EPSCs and IPSCs of the cell shown in **d**, but in the added presence of cholinergic receptor antagonist (50 μΜ hexamethonuim; HEX). **h**, Plot of the E/I offsets during preferred and null direction motion under control conditions (black squares) and in the presence of HEX (red circles). Note that blocking cholinergic transmission leads to a loss of the E/I offsets in the preferred direction, while creating a negative offset (inhibition leads) in the null direction. Data are represented as mean ± SEM.

Since temporal asymmetries are observed under conditions in which photoreceptor/glutamate receptors are blocked, it indicates that they arise from the starburst network itself. Indeed previous studies have provided strong evidence for a differential functional connectivity of cholinergic/GABAergic synapses to DSGCs, whereby inhibitory inputs arise from null-side starbursts while cholinergic inputs arise from both preferred- and null-side starbursts (Chen et al., 2016; Fried et al., 2002; Lee et al., 2010; Yonehara et al., 2011) (null- or preferred-side starbursts refer to those cells with their somas displaced to the side of the DSGC’s receptive field from which null- or preferred-direction stimuli enter, respectively; Fig. 4c). Symmetrical cholinergic receptive fields are presumably mediated by paracrine mechanisms and thus do not reflect the asymmetrical ‘hard-wiring’ (Briggman et al., 2011; Brombas et al., 2017). Thus, the temporal asymmetries in E/I appear to be a natural consequence of the differential connectivity pattern of ACh and GABA synapses from starburst to DSGCs.

**Fig. 4:**
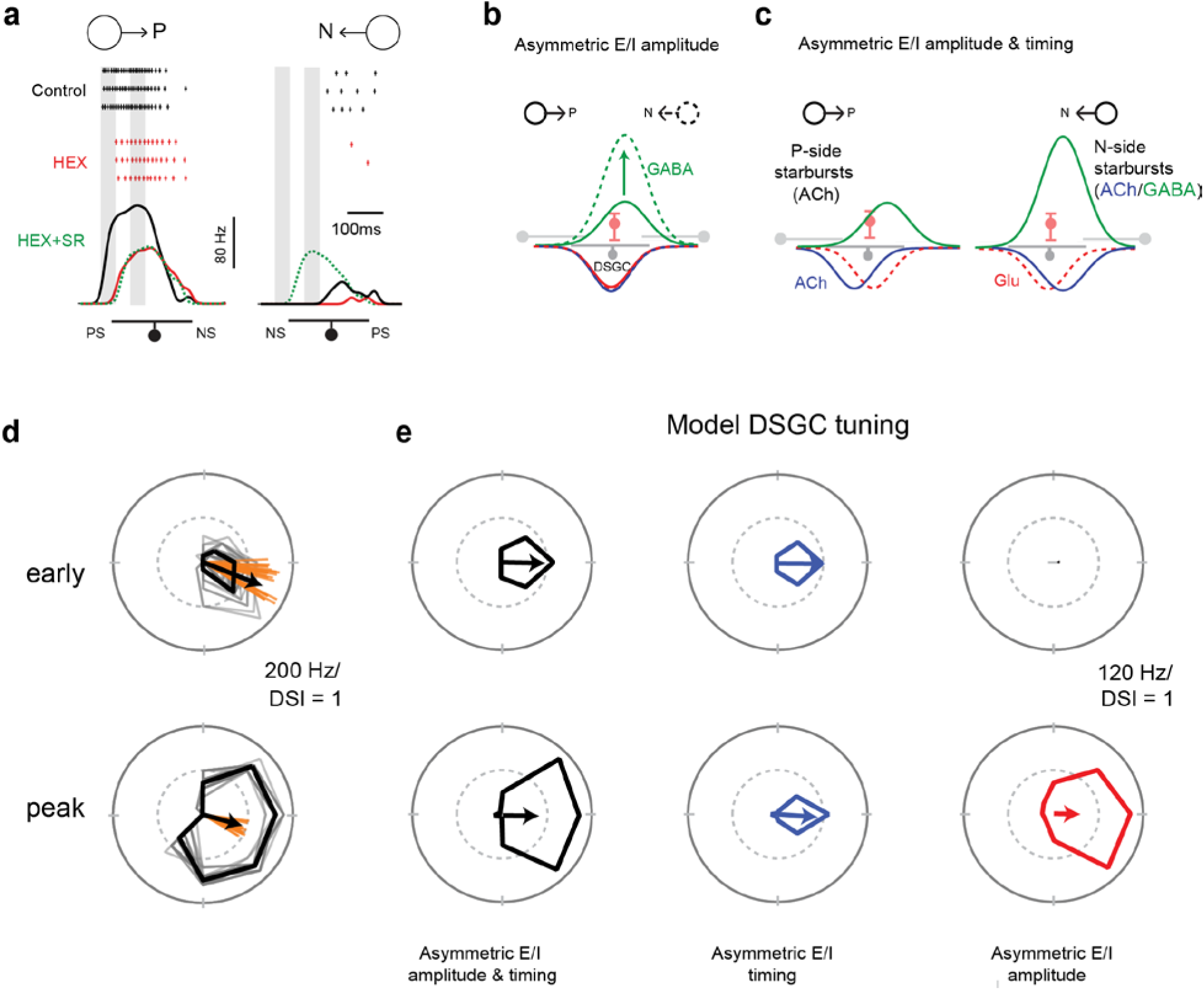
The coordinated actions of two ‘core’ synaptic mechanisms drive direction selectivity in ganglion cells. **a**, Spike rates during <u>stimulus</u> motion in the preferred or null directions in Control (black), HEX (red) to block cholinergic excitation, or HEX+SR-95531 (5 μΜ; green) to isolate the glutamate receptive field in preferred and null directions (PS = preferred starburst side, NS = null starburst side). These effects of HEX on the latency of the response were reversible, and latency was not increased when response amplitudes were decreased using glutamate receptor blockade (see Fig. S1). **b**, Schematic of the current model in which DS is driven largely by changes in the amplitude of inhibition. **c**, DS is driven by asymmetries in E/I temporal offsets and amplitude ratios. In this model, both ACh and GABA are spatially offset, shaping the timing of E/I in a ‘push-pull’ manner. **d**, Polar plots of a DSGC’s early (*top*) and peak (*bottom*) spike responses (the 50 ms shaded regions in a) over 29 trials. The mean response is shown in black, while the individual trials are shown in grey. Spiking during the early phase is sharply tuned then becomes more broadly tuned towards the peak phase of the response. The vector sums of individual trials (orange) indicated that the variance in the early response was higher than the peak response. **e**, As in **d**, but generated using a computational model of a DSGC with both E/I temporal offsets and amplitude asymmetries (*left*, as in **c**), only E/I temporal offsets (*middle*) or only E/I amplitude asymmetry (*right*, as in **b**). E/I offsets alone resulted in a sharper tuning similar to the early responses in **d**, but the tuning becomes broader when E/I ratios are intact, similar to the peak responses in **d** (See Fig. S2 for synaptic currents and spiking responses measured in the model).

Importantly, temporal asymmetries were also observed under natural conditions i.e. when responses were driven by photoreceptors (Fig. 3d-f). Similar to the optogenetic experiments, the magnitude of the E/I offset was greatest for preferred-direction motion and progressively decreased as the stimulus direction approached the DSGC’s null-direction, suggesting that they could also be shaped by offset starburst inputs. Consistent with this notion, blocking nicotinic ACh receptors using a specific antagonist (50 μM hexamethonium) increased the onset latency for excitation evoked by motion in the preferred-direction, and largely abolished the E/I offsets. For null-direction motion, blocking these receptors resulted in a negative offset, as the GABAergic inhibition arising from null-side starbursts were no longer balanced by cholinergic inputs but continued to arrive before glutamate inputs (Fig. 3g). Thus, both ACh and GABA signals arising from starburst dendrites are spatially offset relative to glutamate inputs, unlike previously envisioned (Brombas et al., 2017; Fried et al., 2005; Lee et al., 2010). This effectively creates patterns of ACh/GABA that are temporally synchronized in the null- but not preferred-direction (Fig. 4c), providing the substrate for a timing based mechanism for generating direction selectivity in ganglion cells.

### The temporal evolution of the direction-selective responses in DSGCs

Next, we explored the possible computational benefits offered by the distinct DS mechanisms relying on temporal and amplitude E/I asymmetries. When the spiking responses were averaged over the entire trial, the direction selectivity generated with or without DS inhibition was similar, and thus the DS mechanisms appear redundant (Fig. 2b). However, when the DS tuning was examined on a finer time-scale, it showed distinct characteristics in the early and peak phases of the response, raising the possibility that the two mechanisms operate on distinct time-scales. Specifically, at the onset of the response the DS tuning was sharp but then broadened as the response approached its peak, resulting in a marked decrease in the DSI (early DSI = 0.68 ± 0.06; peak DSI = 0.45 ± 0.06; n = 8; p < 0.01; Fig. 4d). In addition, direction encoded by the initial responses was less reliable than that coded by the peak responses (standard deviation of the vector angle: early = 16 ± 2^0^, peak = 8 ± 2; n = 8; p < 0.005; Fig. 4d). Given that early spikes appear to be driven largely by cholinergic excitation (Fig. 4a, Fig. 3d; but not by glutamate inputs; Fig 4 – Figure Supplement 1), we envisioned that E/I offsets are likely to be more important in determining direction selectivity during the initial phase of the response, while the later phase would likely be dominated by E/I amplitude differences. It is important to note that both DS mechanisms rely on transmitter release from starbursts and thus operate on roughly the same spatial scales. To test these ideas, we next constructed a computational model of the DS circuit in which we could easily control the timing and amplitude of E/I, independently (see Methods for details).

Indeed, the characteristic properties of the DSGC’s direction tuning were reproduced in a model DSGC driven with both temporal and amplitude asymmetries in E/I that arise from the specific arrangement of GABA, ACh and glutamate inputs (Fig. 4c; Fig 4 – Figure Supplement 2 illustrates the synaptic inputs/spiking response of the model DSGC). When starbursts were rendered non-DS in the model DSGC, (Fig. 4c), temporal asymmetries alone were sufficient to generate DS responses with sharp tuning (Fig. 4e), as observed in the ChR2*-Gabra2* KO mouse. When E/I offsets were removed, modulating the peak amplitude of inhibition generated robust DS responses, but with wider tuning. However, in this model lacking offsets, the early responses were lost (Fig. 4e). Thus, the DS mechanisms based on temporal and amplitude asymmetries appear to be complementary, each conferring distinct advantages: the former enables DSGCs to respond in a direction-selective manner sooner than they would have done otherwise, while the latter enables DSGCs to encode direction with higher fidelity albeit on a slower time scale. Interestingly, the shaping of direction selectivity by distinct mechanisms during the early and late response periods is also observed in the primate brain (Pack and Born, 2001; Thiele et al., 2004).

### Conclusions

Our results suggest that two fundamental mechanisms generate direction-selectivity in mouse retina, both originating from unique specializations of the starburst network. The first mechanism relies on temporal asymmetries arising from the asymmetric/symmetric GABAergic/cholinergic connectivity of starburst cells to DSGCs (Chen et al., 2016; Fried et al., 2002; Lee et al., 2010; Yonehara et al., 2011). The second mechanism, which is well established, relies on amplitude asymmetries arising from the DS starburst dendrites. Thus, it appears that the specific connectivity patterns provide the framework over which the specialized properties of starbursts can operate upon to further amplify and fine-tune the direction-selective output of the retina.

The demonstration that the parallel circuit mechanisms driving direction selectivity are individually dispensable necessitates a re-evaluation of the conclusions drawn from a multitude of studies carried out over the last several decades that consider only a single ‘core’ DS mechanism. In the cases of pinpointing the mechanisms generating direction selectivity in starbursts (Ding et al., 2016; Fransen and Borghuis, 2017; Kim et al., 2014; Lee and Zhou, 2006; Munch and Werblin, 2006), or understanding whether timing (Koch et al., 1982; Taylor et al., 2000; Torre and Poggio, 1978) or amplitude ratios generate DS in ganglion cells, the realization of multiple DS mechanisms helps settle divergent views.

However, in other cases, having a second DS mechanism requires a complete re-interpretation of the results. For example, numerous studies over the last forty years have found that blocking cholinergic receptors does not affect direction selectivity in DSGCs and have taken this to mean that ACh does not play an integral role in DS (reviewed by Mauss et al., 2017; Vaney et al., 2012), but rather provides an additional source of non-directional excitation to boost DSGC responses (Brombas et al., 2017; Chen et al., 2016; Lee et al., 2010; Sethuramanujam et al., 2016). In sharp contrast, our results here indicate that in fact ACh signals (unlike glutamate signals) are highly directional, not so much in their amplitudes as previously envisioned (Grzywacz et al., 1998; Grzywacz et al., 1997; Pei et al., 2015), but rather by virtue of their relative timing with inhibition (Fig. 4c). Finally, the array of DS mechanisms described here also raises the interesting issue as to if and how they facilitate the ability for the DSGCs to encode direction over a wide range of stimulus conditions.

## Methods

### Animals

Experiments were performed using adult (either sex) Trhr-EGFP (RRID: MMRRC_030036-UCD) or ChAT-IRES-Cre (RRID: MGI_5475195) crossed with Ai32 (RRID: MGI_5013789) with or without Gabra2^tm22Uru^ (RRID: MGI_5140553). All procedures were performed in accordance with the Canadian Council on Animal Care and approved by the University of Victoria’s Animal Care Committee

### Physiological Recordings

Mice were dark-adapted for approximately 30–60 min before being briefly anesthetized and decapitated. The retina was extracted and dissected in Ringer’s solution under infrared light. The isolated retina was then mounted on a 0.22 mm membrane filter (Millipore) with a pre-cut window to allow light to reach the retina, enabling the preparation to be viewed with infrared light using a Spot RT3 CCD camera (Diagnostic Instruments) attached to an upright Olympus BX51 WI fluorescent microscope outfitted with a 40× water-immersion lens (Olympus Canada). The isolated retina was then perfused with warmed Ringer’s solution (35–37 °C) containing 110 mM NaCl, 2.5 mM KCl, 1 mM CaCl_2_, 1. 6 mM MgCl_2_, 10 mM dextrose and 22 mM NaHCO_3_ that was bubbled with carbogen (95% O_2_:5% CO_2_).

DSGCs were identified by their genetic labeling or by their characteristic DS responses. Light stimuli, produced using a digital light projector (Hitachi Cpx1, refresh rate 75 Hz), were focused onto the outer segments of the photoreceptors using the sub-stage condenser. The background luminance, measured with a calibrated spectrophotometer (Ocean Optics), was set to ~10 photoisomerisations/s (R*/sec). Visual stimuli created in the Matlab environment (Psychtoolbox) were of positive contrasts, ranging between 15% and 1,000% (Weber contrast). Stimulus intensity was increased by 5 log units using neutral density filters to stimulate SAC-ChR2. Light-evoked activity was measured for 200 μm spot moving in eight directions at 1–1.6 mm/s.

Spike recordings were made using the loose cell-attached patch-clamp technique using 5–10-MΩ electrodes filled with Ringer’s solution. Voltage-clamp whole-cell recordings were made using 4–7-MΩ electrodes containing 112. 5 mM CH3CsO3S, 7. 75mM CsCl, 1 mM MgSO4, 10 mM EGTA, 10 mM HEPES, 5 mM QX-314 (Tocris) and 100 μM spermine (Abcam Biochemicals). The pH was adjusted to 7.4 with CsOH. The reversal potential for chloride was calculated to be – 56 mV. The junction potential for the intracellular solution was measured as −8 mV and was corrected offline. Recordings were made with a MultiClamp 700B amplifier (Molecular Devices). Signals were digitized at 10 kHz (PCI-6036E acquisition board, National 9 Instruments) and acquired using custom software written in LabVIEW. Unless otherwise noted, all reagents were purchased from Sigma-Aldrich Canada. D-AP5, and UBP310 were purchased from ABCAM Biochemicals. DL-AP4, SR-95531 and CNQX were purchased from Tocris.

### Computational Modeling

A detailed multi-compartmental model was built using the NEURON simulation environment (Hines and Carnevale, 1997) based on a morphological reconstruction of a real DSGC. Synapses on terminal dendrites throughout the dendritic arbor house glutamatergic (AMPARs), cholinergic (ACH) and GABAergic inputs, whose spatio-temporal offsets and probabilities of release could be varied to match experimental data. Synaptic inputs were activated by a simulated bar moving over the model DSGC in 8 directions at 1mm/s. Cholinergic inputs were symmetrical across the simulated directions, with their probabilities of release (Pr) set to 0.5 and their spatial offsets similarly consistent (50 μm; 50ms at 1mm/s). For control conditions, the inhibitory inputs were highly asymmetric, scaling down from ~0.5 Pr in the null, to ~.012 in the preferred direction, while their spatial offsets also decreased (from ~50 μm in the null to ~0 μm in the preferred). By contrast AMPA receptor-mediated conductances were modelled to occur with no offset relative to their position on the DSGC’s dendritic arbour, simply being activated on average when the simulated light stimulus passed over their location.

### Quantification and statistical analysis

DSI was calculated as (Taylor and Vaney 2002):

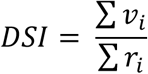

where v_i_ are vectors pointing in the direction of the stimulus and having length r_i_, equal to the number of spikes recorded during that stimulus. DSI ranged from 0 to 1, with 0 indicating a perfectly symmetrical response, and 1 indicating a response in only one of eight directions. The angle was calculated from the direction of the resultant of Σv_i_.

First, the 20–80% rise time of the synaptic currents (EPSCs or IPSCs) was fit by a straight line. The response onset latency was measured as the point at which the extrapolated linear fit crossed the x axis (time axis). The E/I offsets were calculated as difference in the excitatory and inhibitory onsets. Positive offsets indicate that excitation leads inhibition and vice versa. For the purposes of estimating DSI, negative offsets were set to 0.

The spike trains in control conditions were aligned to the edge of the glutamate receptive field, measured either by the spike activity or EPSCs in glutamate isolation i.e. in cholinergic and GABA receptor antagonists (Fig. 4a). After alignment, the number of spikes occurring before the stimulus entered the glutamate receptive field (~50ms in the preferred direction) was considered as the early phase responses. These spikes were completely blocked by cholinergic receptor antagonists (Fig. 4a). The peak phase was estimated as a ~50 ms region close to the peak firing rate in the preferred direction (Fig. 4a).

Population data have been expressed as mean ± SEM and are indicate in the figure legend along with the number of samples. Student’s t test was used to compare values under different conditions and the differences were considered significant when p≤0.05 unless otherwise noted in the figure legend.

